# A history of previous childbirths is linked to women’s white matter brain age in midlife and older age

**DOI:** 10.1101/2020.11.20.391698

**Authors:** Irene Voldsbekk, Claudia Barth, Ivan I. Maximov, Tobias Kaufmann, Dani Beck, Geneviève Richard, Torgeir Moberget, Lars T. Westlye, Ann-Marie G. de Lange

## Abstract

Maternal brain adaptations occur in response to pregnancy, but little is known about how parity impacts white matter (WM) and WM ageing trajectories later in life. Utilising global and regional brain-age prediction based on multi-shell diffusion MRI data, we investigated the association between previous childbirths and WM brain age in 8,895 women in the UK Biobank cohort (age range = 54 - 81 years). The results showed that number of previous childbirths was negatively associated with WM brain age, potentially indicating a protective effect of parity on brain WM later in life. Both global WM and grey matter brain age estimates showed unique contributions to the association with previous childbirths, suggesting partly independent processes. Corpus callosum contributed uniquely to the global WM association with previous childbirths, and showed a stronger relationship relative to several other tracts. While our findings demonstrate a link between reproductive history and brain WM characteristics later in life, longitudinal studies are required to establish causality and determine how parity may influence women’s WM trajectories across the lifespan.

## 1. Introduction

Maternal brain adaptations have been shown during pregnancy and postpartum, with dynamic alterations across brain regions at different time windows since pregnancy [1, 2, 3, 4, 5]. Some of these alterations involve regions implicated in empathy, mentalising, and emotion regulation, and may thus represent adaptations to meet the needs and demands of the offspring, and to secure adequate expression of maternal caregiving [6, 7, 8, 9, 10]. Recent studies also indicate that some effects of pregnancy may be long-lasting [2, 5], potentially influencing brain trajectories later in life [11, 12, 13, 14]. However, neuroimaging studies of the maternal brain have largely focused on grey matter (GM) volume [2, 14, 15, 16, 17, 4, 18] and cortical thickness [3, 19], and less is known about the effects of pregnancy on brain white matter (WM).

Emerging evidence from animal models suggests that pregnancy may induce WM plasticity [20, 21, 22]. Specifically, pregnant mice exhibit increases in oligodendrocyte progenitor cell proliferation, oligodendrocyte generation, and in the number of myelinated axons, indicating an enhanced capacity for myelination in the maternal brain [20]. Pregnancy-induced remyelination may partly explain why pregnancy seem to cause remission of multiple sclerosis (MS), an auto-immune disease that attacks the myelin sheath [23]. In line with this, slower disability progression has been found in parous MS patients after 18 years, compared with nulliparous patients [24]. This effect was strongest in patients that gave birth after disease onset, indicating favourable effects of pregnancy-related adaptations on disease mechanisms in MS.

While the influence of childbirth on WM trajectories in healthy women is largely unknown, one study reported larger regional WM volumes in mothers compared to non-mothers, as well as maternal WM increases that were linked to changes in empathetic abilities during the postpartum period [18]. In line with these findings, a diffusion tensor imaging (DTI) [25] study in rats found that fractional anisotropy (FA), which quantifies the degree of diffusion directionality, increased significantly in the dentate gyrus during pregnancy. However, whole-brain diffusivity also increased in pregnant rats compared to nulliparous rats [21], indicating global changes in the characteristics of molecular water movement - potentially linked to increased extracellular water in the brain during pregnancy [26].

Recent research assessing longitudinal changes in human brain morphology during pregnancy found no WM changes in mothers [2], nor in female adolescents in a follow-up study comparing longitudinal changes in mothers and two years of pubertal development [27]. However, as adolescence is known to involve substantial WM remodelling [28, 29, 30, 31, 32], the lack of effects could possibly reflect insensitivity of the methods used to assess WM changes (T1-weighted estimation of WM volume [2] and gyral WM thickness [27]). In development and ageing studies, WM is commonly investigated using DTI [25], which yields metrics that are highly sensitive to age [33]. However, the accuracy of the DTI approach is limited by factors such as crossing fibres. These obstacles have motivated the development of advanced biophysical diffusion models including white matter tract integrity (WMTI) [34], which is derived from diffusion kurtosis imaging (DKI) [35], and spherical mean technique (SMT) [36, 37]. In contrast to DTI, the DKI model yields metrics estimating the degree of non-Gaussian diffusion, believed to better reflect the complexity of white matter tissue structure [35, 38]. Based on assumptions about the underlying tissue architecture, the WMTI and SMT models enable estimation of the separable contribution of diffusion in the intra- and extra-axonal space [39], which may provide higher biological specificity, i.e. additional information about the microstructural environment [38]. However, the WMTI model does not consider the non-straight and non-parallel nature of fibre crossings and orientation dispersion, something that is factored out in the SMT model to overcome this limitation [36, 37].

In the current study, we utilised four diffusion models (DTI, DKI, WMTI, SMT) to predict WM brain age, and investigated associations between brain-age estimates and previous childbirths in a sample of 8,895 UK Biobank women (mean age *±* standard deviation = 62.45 *±* 7.26). In line with studies suggesting that distinct and regional brain-age estimates may provide additional detail [15, 40, 41, 42], we estimated i) global WM brain age, ii) global GM brain age to test for modality-specific contributions, and iii) WM brain age in 12 major WM tracts in order to identify regions of particular importance.

## 2. Methods and Materials

### 2.1. Sample characteristics

The initial sample was drawn from the UK Biobank (www.ukbiobank.ac.uk), and included 9,899 women. 899 participants with known brain disorders were excluded based on ICD10 diagnoses (chapter V and VI, field F; *mental and behavioural disorders*, including F00 - F03 for Alzheimer’s disease and dementia, and F06.7 *‘Mild cognitive disorder’*, and field G; *diseases of the nervous system*, including inflammatory and neurodegenerative diseases (except G55-59; *“Nerve, nerve root and plexus disorders”*). An overview of the diagnoses is provided in the UK Biobank online resources (http://biobank.ndph.ox.ac.uk/showcase/field.cgi?id=41270), and the diagnostic criteria are listed in the ICD10 diagnostic manual (https://www.who.int/classifications/icd/icdonlineversions). In addition, 99 participants were excluded based on magnetic resonance imaging (MRI) outliers (see section 2.2) and 11 participants were excluded based on missing data on the number of previous childbirths, yielding a total of 8,895 participants that were included in the study. Sample demographics are provided in Table 1.

**Table 1:** Sample demographics. For variables with missing data, sample size (N) is indicated in parentheses. SD = Standard deviation, GCSE = General Certificate of Secondary Education, NVQ = National Vocational Qualification.

|  |  |
| --- | --- |
| **Total N** | 8,895 |
| **Age** |  |
| Mean ± SD | 62.40 ± 7.25 |
| Range [years] | 45.13 - 80.66 |
| **Number of childbirths (live)** |  |
| Mean ± SD | 1.74 ± 1.15 |
| Range | 0 - 8 |
| N in each group:<br>**0** = 1,825 — **1** = 1,190 — **2** = 3,911<br>**3** = 1,535 — **4** = 348 — **5** = 55<br>**6** = 26 — **7** = 4 — **8** = 1 |  |
| **Age at first birth (N = 7,066)** |  |
| Mean ± SD | 26.82 ± 4.99 |
| Range | 14 - 47 |
| **Years since last birth (N = 5,875)** |  |
| Mean ± SD | 32.41 ± 9.21 |
| Range | 6.77 - 55.19 |
| **Menopausal status (N = 8,888)** |  |
| *Yes* | 2,745 |
| *No* | 4,767 |
| *Not sure, had hysterectomy* | 925 |
| *Not sure, other reason* | 451 |
| **Ethnic background (N = 8,872)** |  |
| % White | 97.59 |
| % Black | 0.54 |
| % Mixed | 0.50 |
| % Asian | 0.62 |
| % Chinese | 0.35 |
| % Other | 0.38 |
| % Do not know | 0.02 |
| **Education (N = 8,868)** |  |
| % University/college degree | 42.04 |
| % A levels or equivalent | 13.97 |
| % O levels/GCSE or equivalent | 22.62 |
| % NVQ or equivalent | 3.23 |
| % Professional qualification | 5.79 |
| % None of the above | 6.47 |
| **Assessment location (imaging)** |  |
| Newcastle | 1,419 |
| Cheadle | 7,476 |

### 2.2. MRI data acquisition and processing

A detailed overview of the UK Biobank data acquisition and protocols is available in [43] and [44]. For the diffusion-weighted MRI data, a conventional Stejskal-Tanner monopolar spin-echo echo-planar imaging sequence was used with multiband factor 3. Diffusion weightings were 1000 and 2000 s/mm^2^ and 50 non-coplanar diffusion directions per each diffusion shell. The spatial resolution was 2 mm^3^ isotropic, and 5 anterior-posterior vs 3 anterior-posterior images with b = 0 s/mm^2^ were acquired. All diffusion data were processed using an optimised diffusion pipeline [45] consisting of 6 steps: noise correction [46, 47], Gibbs-ringing correction [48], estimation of echo-planar imaging distortions, motion, eddy-current and susceptibility-induced distortion corrections [49, 50], spatial smoothing using fslmaths from FSL (version 6.0.1) [51] with a Gaussian kernel of 1mm^3^, and diffusion metrics estimation. DTI and DKI derived metrics were estimated using Matlab R2017a (MathWorks, Natick, Massachusetts, USA) as proposed by Veraart and colleagues [52]. The DTI metrics included mean diffusivity (MD), FA, axial diffusivity (AD), and radial diffusivity (RD) [25].

The DKI metrics included mean kurtosis (MK), axial kurtosis (AK), and radial kurtosis (RK) [35]. WMTI metrics included axonal water fraction (AWF), extra-axonal axial diffusivity (axEAD), and extra-axonal radial diffusivity (radEAD) [34]. SMT metrics included intra-neurite volume fraction (INVF), extra-neurite mean diffusivity (exMD), and extraneurite radial diffusivity (exRD) [36]. See [45] for details on the processing pipeline.

Tract-based spatial statistics (TBSS) was used to extract diffusion metrics in WM [53]. Initially, all maps were aligned to the FMRIB58 FA template supplied by FSL, using non-linear transformation in FNIRT [54]. Next, a mean FA image of 18,600 UK Biobank subjects was obtained and thinned to create a mean FA skeleton. The number N = 18,600 was obtained from the processing of the two first UKB data releases. The maximal FA values for each subject were then projected onto the skeleton to minimise confounding effects due to partial volumes and any residual misalignments. Finally, all diffusion metrics were projected onto the subject-specific skeletons. WM features were extracted based on John Hopkins University (JHU) atlases for white matter tracts and labels (with 0 thresholding) [55], yielding a total of 910 WM features including mean values and regional measures for each of the diffusion model metrics. For the region-specific brain age models, 12 tracts of interest used in previous ageing and development studies were extracted [56, 57]; anterior thalamic radiation (ATR), corticospinal tract (CST) cingulate gyrus (CG), cingulum hippocampus (CING), forceps major (FMAJ), forceps minor (FMIN), inferior fronto-occipital fasciculus (IFOF), inferior longitudinal fasciculus (ILF) superior longitudinal fasciculus (SLF), uncinate fasciculus (UF), superior longitudinal fasciculus temporal (SLFT), and corpus callosum (CC). The diffusion MRI data passed TBSS post-processing quality control using the YTTRIUM algorithm [58], and were residualised with respect to scanning site using linear models.

For the GM data, raw T1-weighted MRI data for all participants were processed using a harmonised analysis pipeline, including automated surface-based morphometry and subcortical segmentation. In line with recent brain-age studies [14, 41, 59, 60], we utilised a fine-grained cortical parcellation scheme [61] to extract cortical thickness, area, and volume for 180 regions of interest per hemisphere, in addition to the classic set of subcortical and cortical summary statistics from FreeSurfer (version 5.3) [62]. This yielded a total set of 1,118 structural brain imaging features (360/360/360/38 for cortical thickness/area/volume, as well as cerebellar/subcortical and cortical summary statistics, respectively). Linear models were used to residualise the T1-weighted MRI data with respect to scanning site, intracranial volume [63], and data quality using Euler numbers [64] extracted from FreeSurfer. To remove poor-quality data likely due to motion, participants with Euler numbers of standard deviation (SD) *±* 4 were identified and excluded (n = 80). In addition, participants with SD *±* 4 on the global MRI measures mean FA, mean cortical GM volume, and/or subcortical GM volume were excluded (n = 10, n = 5 and n = 4, respectively), yielding a total of 8,895 participants with both WM (diffusion-weighted) and GM (T1-weighted) MRI data.

### 2.3. Brain-age prediction

Brain-age prediction is a method in which a machine learning algorithm estimates an individual’s age based on their brain characteristics [65]. This estimation is then compared to the individual’s chronological age to estimate each individual’s brain-age gap (BAG), which is used to identify degrees of deviation from normative ageing trajectories. Such deviations have been associated with a range of clinical risk factors [42, 60, 66] as well as neurological and neuropsychiatric diseases [67, 68, 69, 41]. They have also been assessed in previous studies of parity and brain age [13, 14, 15, 59].

Separate brain-age prediction models were run for global WM and GM, and for each of the WM tracts using the *XGBoost regressor model*, which is based on a decision-tree ensemble algorithm (https://xgboost.readthedocs.io/en/latest/python). XGBoost includes advanced regularisation to reduce overfitting [70], and uses a gradient boosting framework where the final model is based on a collection of individual models (https://github.com/dmlc/xgboost). For the global WM and GM models, principal component analyses (PCA) were run on the features to reduce computational time. The top 200 PCA components, explaining 97.84% of the total variance, were used as input for the WM model, and the top 700 components, explaining 98.07% of the variance, were used as input for the GM model. The model parameters were set to *maximum depth* = 4, *number of estimators* = 140, and *learning rate* = 0.1 for the the global and tract-specific WM models, and *maximum depth* = 5, *number of estimators* = 140, and *learning rate* = 0.1 for the global GM model, based on randomised searches with 10 folds and 10 iterations for hyper-parameter optimisation.

The models were run using 10-fold cross-validation, which splits the sample into subsets (folds) and trains the model on all subsets but one, which is used for evaluation. The process is repeated ten times with a different subset reserved for evaluation each time. Predicted age estimates for each participant were derived using the Scikit-learn library (https://scikit-learn.org), and BAG values were calculated using (predicted *−* chronological age). To validate the models, the 10-fold cross validations were repeated ten times, and average R^2^, root mean square error (RMSE), and mean absolute error (MAE) were calculated across folds and repetitions.

### 2.4. Statistical analyses

The statistical analyses were conducted using Python 3.7.6. All variables were standardised (subtracting the mean and dividing by the SD) before entered into the analyses, and *p*-values were corrected for multiple comparisons using false discovery rate (FDR) correction [71]. Chronological age was included as a covariate in all analyses, adjusting for age-bias in the brain age predictions as well as age-dependence in number of childbirths [72, 73].

#### 2.4.1. Previous childbirths and global WM brain age

To investigate associations between number of previous childbirths and global WM brain age, a linear regression analysis was run using *global WM BAG* as the dependent variable, and *number of childbirths* as the independent variable. To control for potential confounding factors, the analysis was rerun including variables known to influence brain structure in ageing or number of childbirths; assessment location [74], education [75, 76], IQ (fluid intelligence) [76], ethnic background [77], body mass index (BMI) [75], diabetic status [78, 60], hypertension [78, 60], smoking and alcohol intake [78, 60], menopausal status (’yes’, ’no’, ’not sure, had hysterectomy’, and ’not sure, other reason’) [79, 80], oral contraceptive (OC) [81, 82] and hormonal replacement therapy (HRT) status (previous or current user vs never used) [83, 84, 81], and experience with stillbirth, miscarriage, or pregnancy termination (’yes’, ’no’) [85, 86] as covariates. In total, 6977 women had data on all variables and were included in these analyses. To test for potential non-linear associations, we added number of childbirths squared as an additional independent variable to the previously defined multiple linear regression model. In addition, we tested for differences in WM BAG by number of childbirths by fitting another linear regression model with WM BAG as the dependent variable and number of childbirths (0,1,2,3,4,5-8) as a fixed factor instead of continuous variable (adjusting for age). Women with zero childbirths served as the reference group. Women with 5-8 childbirths were merged due to low numbers in each group (5 = 55, 6 = 26, 7 = 4, 8 = 1). Cohen’s *d* effect sizes [87] were estimated for each comparison.

#### 2.4.2. Previous childbirths and WM versus GM brain age

To compare the contributions of global WM and GM brain age to the association with previous childbirths, a multiple regression analysis was run with both WM and GM based *BAG estimates* as independent variables and *number of childbirths* as the dependent variable, before eliminating one modality at a time to compare the log-likelihood of the full and reduced models. The significance of model differences was calculated using Wilk’s theorem [88] as 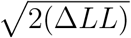, where Δ*LL* = *LL*_1_ *− LL*_2_; the difference in log-likelihood between the reduced model (*LL*_1_) and the full model (*LL*_2_).

Next, we tested for differences between the GM and WM BAG associations with number of previous childbirths using a *Z* test for correlated samples [89] with

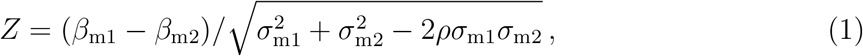

where m1 = model 1 (WM); m2 = model 2 (GM); *β* = beta coefficients from the linear regressions between number of childbirths and each model; *σ* = standard errors of the beta coefficients; *ρ* = age-adjusted correlation between the modality-specific brain age gap estimates. As a follow-up, we tested the associations between previous childbirths and mean FA, mean MD, and total GM volume.

#### 2.4.3. Previous childbirths and regional WM tracts

To test for unique contributions by each tract to the global WM association with previous childbirths, a multiple regression analysis was run with all tract-specific *BAG estimates* as independent variables and *number of childbirths* as the dependent variable, before eliminating the tracts one at a time to compare the log-likelihood of the full and reduced models. The significance of model differences was calculated using Wilk’s theorem as described in Section 2.4.2. In addition, the reduced *χ*^2^ values for each of the models were calculated to account for the difference in number of input variables to the full and reduced models (13 for the full model including 12 tracts + age, versus 11 for the reduced models where each of the tracts were eliminated one by one). Next, we performed separate regression analyses for each tract-specific BAG estimate versus number of childbirths, before testing for differences between the associations using pairwise *Z* tests for correlated samples (Eq. 1; Section 2.4.2).

## 3. Results

The age prediction accuracies for the global WM and GM models, as well as each of the tract-specific WM models are shown in Table 2. The correlations between predicted and chronological age for the global models are shown in Supplementary Information (SI) Figure 1. The associations between number of previous childbirths and BAG estimates based on each of the predictions are shown in Table 3.

**Table 2:** Average R^2^, root mean square error (RMSE), mean absolute error (MAE), and correlation (*r*) between predicted and chronological age for the age prediction models. WM = white matter, GM = grey matter; ATR = anterior thalamic radiation; CST = corticospinal tract; CG = cingulate gyrus; CING = cingulum hippocampus; FMAJ = forceps major; FMIN = forceps minor; IFOF = inferior fronto-occipital fasciculus; ILF = inferior longitudinal fasciculus; SLF = superior longitudinal fasciculus; UF = uncinate fasciculus; SLFT = superior longitudinal fasciculus temporal; CC = corpus callosum; CI = confidence interval.

| Global predictions |  |  |  |  |  |
| --- | --- | --- | --- | --- | --- |
| Modality | $R^2$ | RMSE | MAE | $r$ [95% CI] | $p$ |
| WM | $0.51 \pm 0.02$ | $5.06 \pm 0.11$ | $4.10 \pm 0.09$ | 0.72 [0.71, 0.73] | <0.001 |
| GM | $0.32 \pm 0.02$ | $5.98 \pm 0.13$ | $4.97 \pm 0.11$ | 0.57 [0.55, 0.58] | <0.001 |
| Predictions for each WM tract |  |  |  |  |  |
| Tract | $R^2$ | RMSE | MAE | $r$ [95% CI] | $p$ |
| ATR | $0.31 \pm 0.02$ | $6.03 \pm 0.13$ | $4.92 \pm 0.12$ | 0.56 [0.54, 0.57] | <0.001 |
| CST | $0.15 \pm 0.02$ | $6.69 \pm 0.14$ | $5.53 \pm 0.13$ | 0.38 [0.37, 0.40] | <0.001 |
| CG | $0.19 \pm 0.02$ | $6.54 \pm 0.13$ | $5.38 \pm 0.12$ | 0.44 [0.42, 0.45] | <0.001 |
| CING | $0.12 \pm 0.02$ | $6.81 \pm 0.14$ | $5.64 \pm 0.12$ | 0.34 [0.32, 0.36] | <0.001 |
| FMAJ | $0.14 \pm 0.02$ | $6.71 \pm 0.13$ | $5.55 \pm 0.12$ | 0.38 [0.37, 0.41] | <0.001 |
| FMIN | $0.26 \pm 0.02$ | $6.24 \pm 0.13$ | $5.09 \pm 0.12$ | 0.51 [0.49, 0.52] | <0.001 |
| IFOF | $0.25 \pm 0.02$ | $6.29 \pm 0.13$ | $5.16 \pm 0.12$ | 0.50 [0.48, 0.51] | <0.001 |
| ILF | $0.18 \pm 0.02$ | $6.55 \pm 0.14$ | $5.40 \pm 0.13$ | 0.43 [0.41, 0.44] | <0.001 |
| SLF | $0.18 \pm 0.02$ | $6.54 \pm 0.13$ | $5.40 \pm 0.12$ | 0.43 [0.41, 0.45] | <0.001 |
| UF | $0.18 \pm 0.03$ | $6.56 \pm 0.13$ | $5.42 \pm 0.12$ | 0.42 [0.40, 0.44] | <0.001 |
| SLFT | $0.17 \pm 0.02$ | $6.58 \pm 0.14$ | $5.42 \pm 0.13$ | 0.42 [0.41, 0.44] | <0.001 |
| CC | $0.25 \pm 0.02$ | $6.26 \pm 0.13$ | $5.13 \pm 0.12$ | 0.50 [0.49, 0.52] | <0.001 |

**Table 3:** Associations between each of the brain age gap estimates and number of previous childbirths (*β*_*CB*_, standard error (SE), *t, p*, and *p*_*corr*_). Chronological age was included in the analyses for covariate purposes and *p*-values are reported before and after correction for multiple comparisons, with corrected *p*-values *<* 0.05 highlighted in bold. WM = white matter, GM = grey matter; ATR = anterior thalamic radiation; CST = corticospinal tract; CG = cingulate gyrus; CING = cingulum hippocampus; FMAJ = forceps major; FMIN = forceps minor; IFOF = inferior fronto-occipital fasciculus; ILF = inferior longitudinal fasciculus; SLF = superior longitudinal fasciculus; UF = uncinate fasciculus; SLFT = superior longitudinal fasciculus temporal; CC = corpus callosum.

| Global associations |  |  |  |  |  |
| --- | --- | --- | --- | --- | --- |
| Modality | $\beta_{CB}$ | SE | $t$ | $p$ | $p_{corr}$ |
| WM | -0.037 | 0.007 | -5.44 | $5.46 \times 10^{-8}$ | **$2.31 \times 10^{-7}$** |
| GM | -0.029 | 0.005 | -5.41 | $6.43 \times 10^{-8}$ | **$2.31 \times 10^{-7}$** |
| Associations for each WM tract |  |  |  |  |  |
| Tract | $\beta_{CB}$ | SE | $t$ | $p$ | $p_{corr}$ |
| ATR | -0.022 | 0.006 | -3.66 | $2.51 \times 10^{-4}$ | **$5.03 \times 10^{-4}$** |
| CST | -0.006 | 0.004 | -1.44 | 0.15 | 0.16 |
| CG | -0.013 | 0.005 | -2.73 | 0.01 | **0.01** |
| CING | -0.013 | 0.004 | -3.17 | 0.00 | **0.00** |
| FMAJ | -0.009 | 0.005 | -2.02 | 0.04 | 0.06 |
| FMIN | -0.021 | 0.006 | -3.73 | $1.90 \times 10^{-4}$ | **$4.26 \times 10^{-4}$** |
| IFOF | -0.012 | 0.006 | -2.24 | 0.03 | **0.04** |
| ILF | -0.008 | 0.005 | -1.59 | 0.11 | 0.14 |
| SLF | -0.006 | 0.005 | -1.18 | 0.24 | 0.24 |
| UF | -0.007 | 0.005 | -1.46 | 0.14 | 0.16 |
| SLFT | -0.011 | 0.005 | -2.15 | 0.03 | **0.04** |
| CC | -0.029 | 0.006 | -5.25 | $1.60 \times 10^{-7}$ | **$4.81 \times 10^{-7}$** |

**Figure 1:**
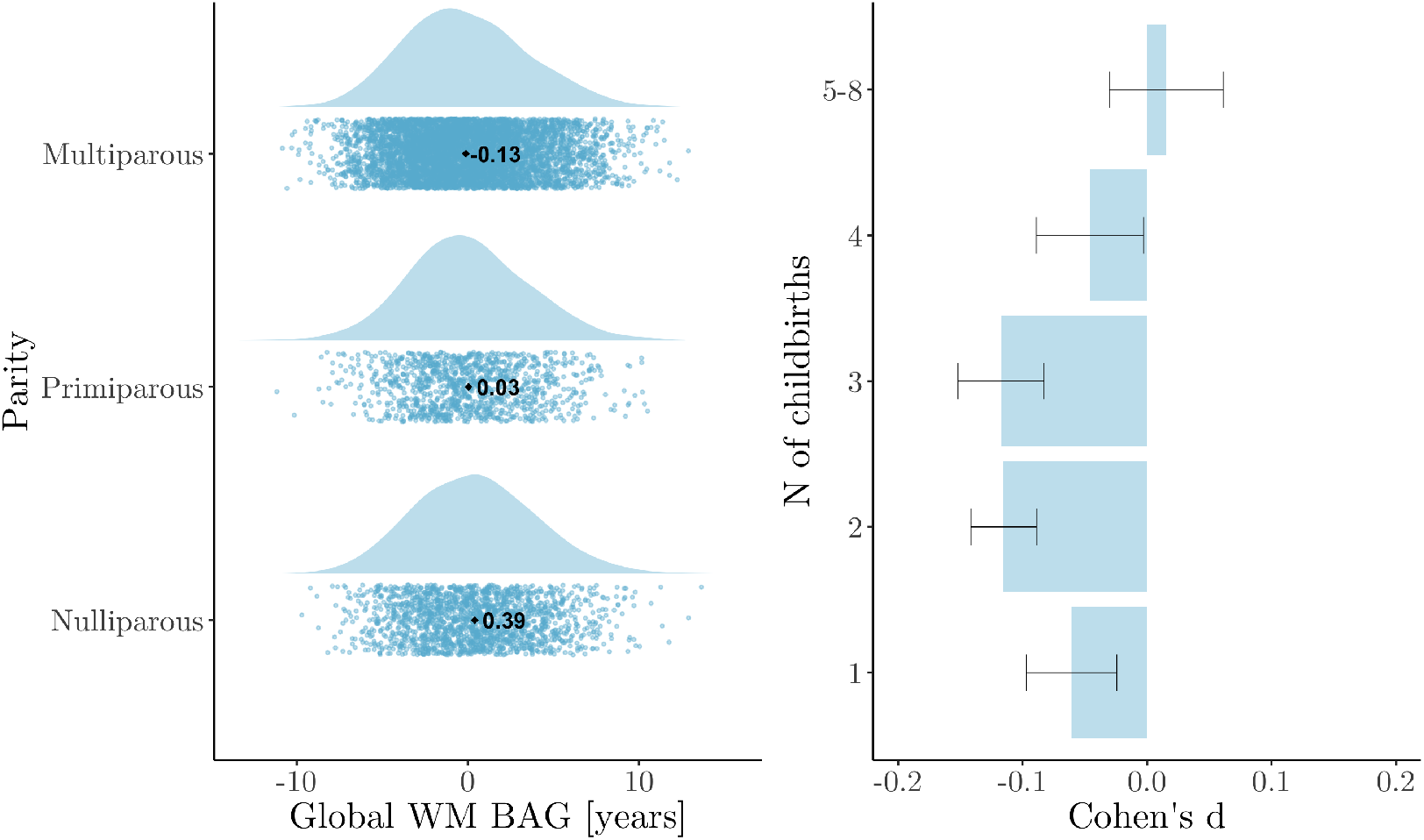
Global white matter (WM) brain age gap (BAG) for groups of women based on number of previous childbirths. **Left plot**: WM BAG in nulliparous, primiparous, and multiparous women are displayed as raincloud plots, which combines raw data points (scatterplot) and the distributions of the data (histogram) using split-half violins. The mean for each group is displayed as red dot and text. **Right plot**: Cohen’s *d* effect sizes for differences between each group of parous women (1,2,3,4,5-8) versus nulliparous women.

### 3.1. Previous childbirths and global WM brain age

Global WM BAG showed a negative association with number of previous childbirths, indicating a younger-looking brain in parous women (see Table 3). As shown in Figure 1, mean WM BAG was positive in nulliparous and primiparous women (0.39 and 0.03, respectively) and negative in multiparous women (−0.13). The model including potential confounding factors showed a corresponding association of *β* = *−*0.030, SE = 0.007, *t* = *−*4.06, *p* = 4.94 *×* 10^*−*5^, indicating that assessment location, education, IQ, ethnic background, BMI, diabetic status, hypertension, smoking and alcohol intake, menopausal status, and OC and HRT use could not fully explain the association between number of childbirths and global WM BAG. The correlations between global WM BAG and demographics, covariates, and number of childbirths are shown in SI Figure 2. Number of previous childbirths and age at first birth correlated *r* = *−*0.30, *p* = 2.82 *×* 10^*−*138^ (adjusted for age). To test for an association with global WM brain age, an analysis was run with WM BAG as the dependent variable and *age at first birth* as independent variable, including all covariates (assessment location, education, IQ, ethnic background, BMI, diabetic status, hypertension, smoking and alcohol intake, menopausal status, and OC and HRT use). No association was found (*β* = *−*0.002, SE = 0.009, *t* = *−*0.27, *p* = 0.79, N = 5515). In addition to a linear effect, we also found evidence for a non-linear association between number of childbirths and global WM BAG: *β* = 0.015, SE = 0.004, *t* = 3.41, *p* = 0.001. Differences in WM BAG were found between nulliparous women and women with one, two, or three previous childbirths. The differences were not significant for women with four or more childbirths (shown in Figure 1 and Table 4).

**Table 4:** Differences in global white matter (WM) brain age gap (BAG) by number of childbirths (*β*, standard error (SE), *t, p*, and Cohen’s *d ±* SE), based on a regression model with WM BAG as the dependent variable and number of childbirths as a fixed factor, where 0 childbirths served as the reference group. Chronological age was included in the analyses for covariate purposes.

| Childbirths | $\beta$ | SE | $t$ | $p$ | $d$ | $N$ |
| --- | --- | --- | --- | --- | --- | --- |
| 0 (intercept) | 0.118 | 0.327 | 0.36 | 0.72 | - | 1825 |
| 1 | -0.368 | 0.132 | -2.80 | $5.14 \times 10^{-3}$ | $-0.06 \pm 0.04$ | 1190 |
| 2 | -0.510 | 0.101 | -5.06 | $4.35 \times 10^{-7}$ | $-0.12 \pm 0.03$ | 3911 |
| 3 | -0.680 | 0.123 | -5.51 | $3.72 \times 10^{-8}$ | $-0.12 \pm 0.02$ | 1535 |
| 4 | -0.330 | 0.208 | -1.59 | 0.11 | $-0.05 \pm 0.04$ | 348 |
| 5-8 | 0.117 | 0.390 | 0.30 | 0.76 | $-0.02 \pm 0.05$ | 86 |

**Figure 2:**
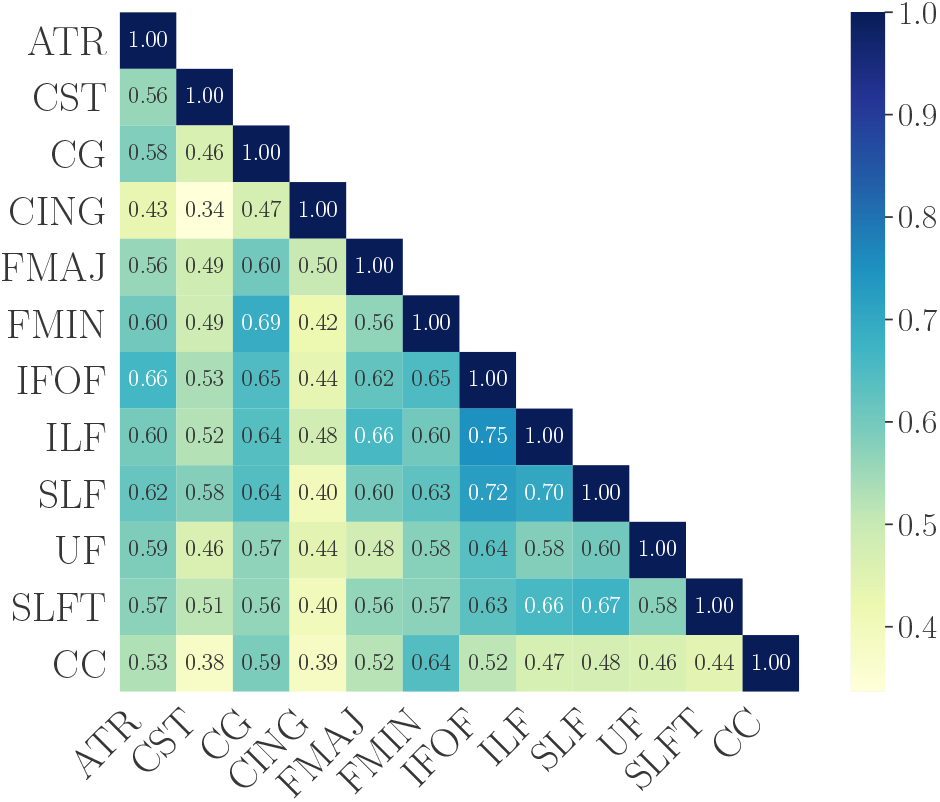
The correlations (Pearson’s *r*) between tract-specific brain age gap (BAG) estimates. The BAG values were first corrected for chronological age using linear models [72], and the residuals were used in the correlation analysis. ATR = anterior thalamic radiation; CST = corticospinal tract; CG = cingulate gyrus; CING = cingulum hippocampus; FMAJ = forceps major; FMIN = forceps minor; IFOF = inferior fronto-occipital fasciculus; ILF = inferior longitudinal fasciculus; SLF = superior longitudinal fasciculus; UF = uncinate fasciculus; SLFT = superior longitudinal fasciculus temporal; CC = corpus callosum.

### 3.2. Previous childbirths and WM versus GM brain age

The age prediction based on the WM model showed higher accuracy compared to the GM prediction (R^2^ of 0.51 versus 0.32), as shown in Table 3. To directly compare the model predictions, a post-hoc *Z* test for correlated samples (Eq. 1; Section 2.4.2) was run on the model-specific fits of predicted versus chronological age (Pearson’s *r* values). The result showed a significant difference in model performance in favour of the WM model; *Z* = - 11.90, *p* = 1.06 *×* 10^*−*32^.

When comparing regression models including both WM and GM-based BAG estimates to models including only one of the modalities, both the WM-based and the GM-based estimates were found to contribute uniquely to the association with number of previous childbirths, as shown in Table 5. The Z test for differences in associations (Eq. 1; Section 2.4.2) revealed similar associations between number of childbirths and WM-based versus GM-based BAG estimates, as shown in Table 6. The follow-up tests of mean FA, mean MD, and total GM volume showed positive associations between number of childbirths and mean FA as well as total GM volume, and a negative association with mean MD, as shown in SI Table 1. Only the association with MD was significant after adjusting for multiple comparisons.

**Table 5:**
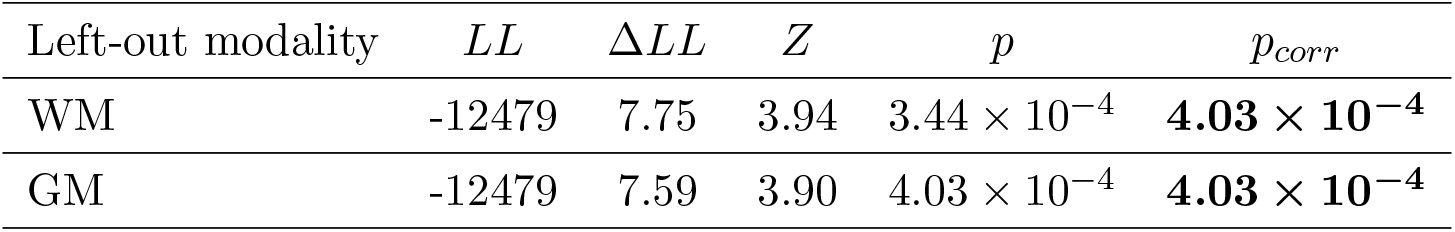
Difference in log-likelihood (Δ*LL*) between regression analyses where grey matter (GM) and white matter (WM)-based brain age gap estimates were eliminated one by one, compared to a model where both were included. The log likelihood (*LL*) value for the model including both modalities was -12471. Reported are *p* values before and after correction for multiple comparisons, with corrected *p*-values *<* 0.05 highlighted in bold.

**Table 6:** Difference in the associations (*β*) between number of previous childbirths and white matter (WM) versus grey matter (GM)-based brain age estimates (Eq. 1). SE = standard error.

| $\beta_{WM} \pm SE$ | $\beta_{GM} \pm SE$ | $Z$ | $p$ |
| --- | --- | --- | --- |
| $-0.037 \pm 0.007$ | $-0.029 \pm 0.005$ | 1.04 | 0.30 |

### 3.3. Previous childbirths and regional WM tracts

Significant (*p <* 0.05) negative associations between number of previous childbirths and WM BAG estimates were found for ATR, CG, CING, FMIN, IFOF, SLFT, and CC, as shown in Table 3. The correlations between the tract-specific BAG estimates are shown in Figure 2. CC contributed uniquely to the global WM association with number of previous childbirths, as shown in Table 7. Pairwise Z tests for differences in associations revealed that ATR and FMIN had significantly stronger associations with previous childbirths compared to SLF, while CC was more strongly associated with previous childbirths than CST, CG, FMAJ, IFOF, ILF, SLF, UF, and SLFT, as shown in Figure 3. As CC showed the most prominent contribution to the association with previous childbirths, we extracted the feature importance ranking from the CC-specific age prediction. WMTI-radEAD showed the highest gain (SI Table 2), indicating that this diffusion metrics was most important for generating the prediction.

**Table 7:** Difference in log-likelihood (Δ*LL*) between regression analyses against *number of previous childbirths* (including age as a covariate). The difference is calculated between models where all tracts are included and models where single tracts are left out one at a time. Reported are *p*-values before and after correction for multiple comparisons, with corrected *p*-values *<* 0.05 highlighted in bold. 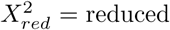 chi-squared values for each reduced model. For the full model, 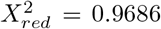. ATR = anterior thalamic radiation; CST = corticospinal tract; CG = cingulate gyrus; CING = cingulum hippocampus; FMAJ = forceps major; FMIN = forceps minor; IFOF = inferior fronto-occipital fasciculus; ILF = inferior longitudinal fasciculus; SLF = superior longitudinal fasciculus; UF = uncinate fasciculus; SLFT = superior longitudinal fasciculus temporal; CC = corpus callosum.

| Left-out tract | $\Delta LL$ | $Z$ | $p$ | $p_{corr}$ | $X_{red}^2$ |
| --- | --- | --- | --- | --- | --- |
| ATR | 1.86 | 1.93 | 0.13 | 0.65 | 0.9689 |
| CST | 0.08 | 0.41 | 0.74 | 0.78 | 0.9685 |
| CG | 0.02 | 0.22 | 0.78 | 0.78 | 0.9685 |
| CING | 1.56 | 1.79 | 0.16 | 0.65 | 0.9688 |
| FMAJ | 0.39 | 0.88 | 0.54 | 0.78 | 0.9686 |
| FMIN | 0.64 | 1.14 | 0.42 | 0.78 | 0.9686 |
| IFOF | 0.02 | 0.19 | 0.78 | 0.78 | 0.9685 |
| ILF | 0.25 | 0.71 | 0.62 | 0.78 | 0.9685 |
| SLF | 1.08 | 1.47 | 0.27 | 0.68 | 0.9687 |
| UF | 1.03 | 1.44 | 0.29 | 0.68 | 0.9687 |
| SLFT | 0.28 | 0.75 | 0.60 | 0.78 | 0.9686 |
| CC | 6.00 | 3.46 | 0.00 | **0.02** | 0.9698 |

**Figure 3:**
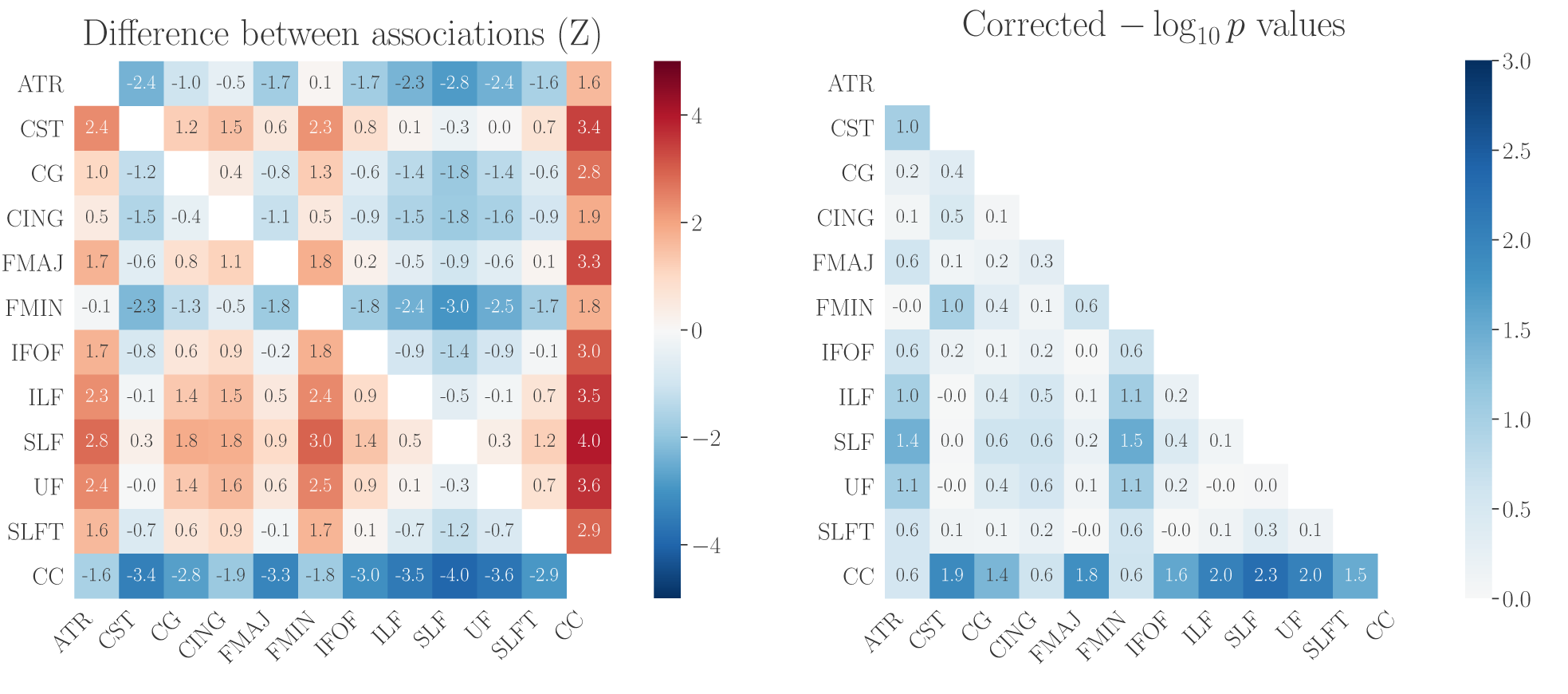
The differences (Z) between tract-specific associations with previous childbirths. Left plot: the values indicate the association with the tract on the y-axis minus the association with the tract on the x-axis (Eq. 1; Section 2.4.2). The beta values for each association are provided in Table 3. Right plot: *p*-values for the differences between associations, reported as the common logarithm (*−log*_10_(*p*)) and corrected for multiple comparisons. A *− log*_10_(*p*) value of *>* 1.3 corresponds to *p <* 0.05. ATR = anterior thalamic radiation; CST = corticospinal tract; CG = cingulate gyrus; CING = cingulum hippocampus; FMAJ = forceps major; FMIN = forceps minor; IFOF = inferior fronto-occipital fasciculus; ILF = inferior longitudinal fasciculus; SLF = superior longitudinal fasciculus; UF = uncinate fasciculus; SLFT = superior longitudinal fasciculus temporal; CC = corpus callosum.

## 4. Discussion

The current study investigated the association between previous childbirths and WM brain age by utilising global and region-specific brain-age prediction. The results showed that a higher number of previous childbirths was associated with lower brain age in global WM, as well as in WM tracts including ATR, CG, CING, FMAJ, FMIN, IFOF, SLFT, and CC. CC contributed uniquely to the global WM association with previous childbirths, and showed a stronger relationship with previous childbirths relative to several other tracts. When assessing global WM compared to GM brain age estimates, both modalities showed unique contributions to the association with previous childbirths. Taken together, these results indicate an association between previous childbirths and global WM ageing later in life, with regional effects that may be particularly prominent in CC.

### 4.1. Previous childbirths and global WM age

During pregnancy, several adaptations in the female body and brain take place in order to meet the needs and demands of the offspring, and to secure adequate expression of maternal caregiving [9, 6]. Maternal adaptation in WM may thus be induced to meet these new demands, by promoting myelination to ensure increased efficiency of neural transmission in relevant WM tracts. While speculative, our results may reflect a long-term benefit of pregnancy-induced WM plasticity, potentially promoting favourable WM trajectories later in life [90]. In support of long-term positive effects of childbirth on WM health, parity is associated with protective effects on age-related decline in learning, memory, and brain health in rats [91]. Further evidence for beneficial effects of parity on brain ageing stems from a study showing that telomeres are significantly elongated in parous relative to nulliparous women [11], suggesting that parity may slow down the pace of cellular ageing.

The current results are also in line with previous studies in MS patients showing beneficial effects of pregnancy on WM health [18, 20, 21, 22, 24]. Oestradiol, a type of oestrogen that increases 300-fold during pregnancy [92], has been linked to pregnancy-induced MS remission [93], likely due to its anti-inflammatory and neuroplastic properties [94]. Further evidence for protective effects of oestradiol stems from hormonal replacement studies in postmenopausal women: long-term oestrogen use has been associated with greater WM volumes [95], indicating a protective effect on WM loss in ageing. Postnatally, oestradiol levels drop rapidly and may promote a pro-inflammatory immune environment [96], which has been linked to a risk of relapse or worsening of symptoms in women suffering from MS [97, 98]. However, in a long-term perspective, pregnancy does not increase the risk of exacerbated disability [99], and some evidence suggests that long-term disability progression improves in MS patients following childbirth [24]. Any influence of pregnancy-related oestrogen fluctuations (i.e. perinatal surge, postpartum drop) on brain ageing is likely to involve a complex interplay of neurobiological processes, and evidence suggests that genetic factors may modulate how oestrogen exposure affects brain health [59, 100, 101]. Beside oestrogen, other hormones such as progesterone, prolactin, oxytocin, and cortisol also fluctuate during pregnancy and may regulate WM plasticity [102, 103, 104]. For instance, emerging evidence from animal models suggests protective effects of progesterone and prolactin on white matter structure due to its pro-myelinating properties [105, 106, 103]. Prolactin-signalling during pregnancy has been linked to increases in oligodendrocyte precursor cells and oligodendrocyte production in the maternal central nervous system, resulting in an enhanced ability to regenerate white matter damage [104]. While the influence of hormone exposure on brain ageing trajectories is currently unclear, other pregnancy-induced adaptations such as the proliferation of regulatory T cells or foetal microchimerism may also represent mechanisms underlying potential long-term benefits of pregnancy on brain ageing (for a review see [102]). Future studies should target the links between hormone- and immune-related neuroplasticity in pregnancy, and the potential effect of these processes on women’s brain ageing trajectories.

Experience-dependent brain plasticity due to parenting is another possible mechanism that may underlie individual differences in WM BAG between parous and nulliparous women. Becoming a parent represents a significant transition in life, including extensive lifestyle changes and brain adaptations in regions relevant for caregiving behaviour to meet the needs and demands of the offspring [6, 7, 8, 9, 10, 96, 97]. For instance, studies have found a link between caregiving behaviour, altered brain activation, and levels of oxytocin in both fathers and mothers [107], and parity has been associated with brain age and cognitive function in both men and women [13]. While experience-dependent brain plasticity related to parenting may influence WM trajectories later in life, animal research has demonstrated that WM adaptations are also induced by pregnancy itself [20, 21, 22]. Hence, pregnancy- and parental experience-induced plasticity are not mutually exclusive, and may together shape WM brain trajectories later in life. To disentangle the effects of pregnancy and parental experience on WM brain ageing trajectories, further research may aim to include fathers as well as women who have experienced adoption (parenting experience without the pregnancy experience) and stillbirth (pregnancy experience without the parenting experience).

While the results showed a negative linear relationship between parity and brain age estimates, follow-up analyses also indicated a quadratic effect in line with what we observed in one of our previous studies based on GM brain age [14]. However, this non-linear GM effect was not replicated in a follow-up study conducted in 8,800 new UK Biobank participants [15], and given the small number of women with *>* 4 children, further studies are needed to conclude on whether any protective effects of parity may be less pronounced in grand-parous women.

Previous childbirths also showed associations with mean FA, mean MD, and total GM volume, but with lower t-values compared to the associations with BAG. Relative to more traditional MRI summary measures, age prediction models have the advantage of encoding normative trajectories of brain differences across age, and condensing a rich variety of brain characteristics into single estimates per individual. Hence, brain age prediction provides a useful summary measure that may serve as a proxy for brain integrity across normative and clinical populations [41, 68, 108, 42, 109].

### 4.2. Modality-specific and regional effects

In line with recent studies demonstrating high age prediction accuracy based on diffusion imaging data [42, 68, 110, 111, 112], the WM prediction showed higher accuracy compared to the GM model, of which the accuracy corresponded to our previous UK Biobank studies [14, 15, 59]. Importantly, we found unique contributions by both models, suggesting that the diffusion-based WM model may pick up variance not explained by the T1-based GM model. These findings highlight the relevance of assessing brain characteristics using different MRI modalities to increase our understanding of possible long-term effects of pregnancy on the brain.

The most prominent regional WM effect of childbirth was seen in the CC, showing both a unique contribution and a stronger association relative to several other tracts, potentially indicating regional variations. While the volume of most WM tracts increase from childhood to young adulthood, peaks around the fifties, and subsequently declines [33, 56, 113, 114, 115, 57], CC volume has been shown to peak already in the beginning of the thirties, exhibiting an earlier onset of age-related decline relative to other WM tracts [57]. Sex differences have also been found in CC ageing, with steeper volumetric decline in men relative to women [116]. Although speculative, our findings could potentially reflect a mitigating effect of parity on age-related CC volumetric decline. While little is known about pregnancy-induced alterations in specific WM regions, an increased number of myelinated axons in the CC have been found in healthy pregnant rats [20], and increased CC remyelination has been observed in pregnant rat models of demyelination [20, 22]. Interestingly, radEAD from the WMTI diffusion model was found to be the most important feature for the CC-specific WM age prediction. WMTI-radEAD has been related to degree of myelination in both *ex vivo* [117] and *in vivo* animal histology models [118], as well as in an *ex vivo* human model of CC [119]. While this may potentially indicate that the CC association with previous childbirths could be driven by individual differences in myelin-related ageing processes, the precise underlying neural substrates of diffusion metrics remain to be clarified. Furthermore, CC is also the most accessible WM structure to investigate given its size and location in the brain, and the relative simple and coherent microstructural milieu may be easier to resolve using diffusion MRI compared to other pathways with more complex tissue structure. The tract extraction procedure could thus result in higher signal-to-noise ratio for the CC than for the remaining tracts, rendering it more sensitive to tests of WM associations with childbirth.

### 4.3. Study limitations

The cross-sectional design of the current study represents a major limitation, and longitudinal studies following women through pregnancy, postpartum, and into midlife and older age are required to infer causality between the observed associations. Furthermore, a complex interplay of numerous underlying processes likely influence the link between parity and WM trajectories. While the current study controls for a range of confounding factors including neurological disease, mental disorders, education, lifestyle behaviours, and cardiovascular risk, the number of children a woman gives birth to — as well as their brain health across the lifespan — may also depend on genetic predispositions, life circumstances, and additional aspects of general health. While information on breastfeeding was not available in the current dataset, this factor is relevant for future studies as it is known to influence estrogen exposure [120] and maternal health [121].

Our results could potentially reflect long-term effects of pregnancy-related processes such as myelination. However, the exact neurobiological underpinnings of diffusion metrics cannot be directly inferred, and although we utilised advanced diffusion modelling which is sensitive to biophysical tissue properties [38], the biological substrates underlying these metrics remain to be elucidated by future studies. In addition, controlling for the effect of extracellular water or indices of hydration [122] as well as including measures of WM hyperintensities [123, 124] could potentially provide more accurate models of WM ageing.

The effect sizes for differences between groups of parous and nulliparous women ranged from 0.06 - 0.12, which is generally considered small. Small effects are common in large datasets [125, 126], and while parity may explain only a small portion of the variance in brain age, our findings emphasise the importance of including female-specific variables in studies of women’s brain ageing, as well as sex differences in risk factors and disease [127]. While the UK Biobank dataset enables detection of subtle effects due to its large sample size, the cohort is homogeneous with regard to ethnic background (97% white participants in the current study), preventing any conclusion about associations between reproductive history and WM ageing across ethnic groups. The cohort is also homogeneous with regard to education level and residing in the United Kingdom, and since access to healthcare, social welfare benefits, and maternity leave policies differ significantly across the world, such factors are important to address in future studies including multiple cohorts. The UK Biobank is also characterised by a “healthy volunteer effect” [128], suggesting that it is not representative of the general population [129]. Hence, the presented results may not apply to populations beyond those represented in this cohort. However, in context of the historical lack of research on women’s brain health [130], the current results may prompt further study into how female-specific factors such as pregnancy influences neural processes involved in normal ageing - as well as autoimmune conditions and Alzheimer’s disease, of which the risks are higher for women relative to men [131, 132].

### 4.4. Conclusion

In summary, the current study found an association between a higher number of previous childbirths and lower WM brain age, in line with previous studies showing relationships between parity and brain characteristics in midlife and older age [13, 15, 19]. As outlined above, a complex interplay of numerous underlying processes likely influence the link between previous childbirths and brain health in older age. Thus, while our results may suggest that reproductive history influences women’s WM ageing trajectories, prospective longitudinal studies assessing this multi-factorial relationship are greatly needed to increase the knowledge about women’s brain health across the lifespan.

## Author contributions

I.V., L.T.W., and A-M.G.dL. designed the study; I.I.M., T.K., C.B., and A-M.G.dL. processed the data; I.V., D.B., and A-M.G.dL. performed the data analyses; I.V., C.B., I.I.M., T.K., D.B., G.R., T.M., L.T.W., and A-M.G.dL. interpreted the data; I.V. and A-M.G.dL. drafted and finalised the manuscript; C.B., I.I.M., T.K., D.B., G.R., T.M., and L.T.W. critically revised the first draft and approved the final manuscript.

## Acknowledgements

This research has been conducted using the UK Biobank under Application 27412. The work was performed on the Service for Sensitive Data (TSD) platform, owned by the University of Oslo, operated and developed by the TSD service group at the University of Oslo IT-Department (USIT). Computations were also performed using resources provided by UNINETT Sigma2 – the National Infrastructure for High Performance Computing and Data Storage in Norway. While working on this study, the authors received funding from the Research Council of Norway (I.V; Student scholarship; T.K.; 276082; L.T.W.; 273345, 249795, 298646, 300768, 223273; C.B.; 250358), the South-East Norway Regional Health Authority (L.T.W.; 2018076, 2019101), the European Research Council under the European Union’s Horizon 2020 research and innovation programme (L.T.W.; 802998), and the Swiss National Science Foundation (AM.G.dL.; PZ00P3 193658).

