## Supplementary Material for "A history of previous childbirths is linked to women’s white matter brain age in midlife and older age"

### Supplementary Information

#### 1. Age prediction models

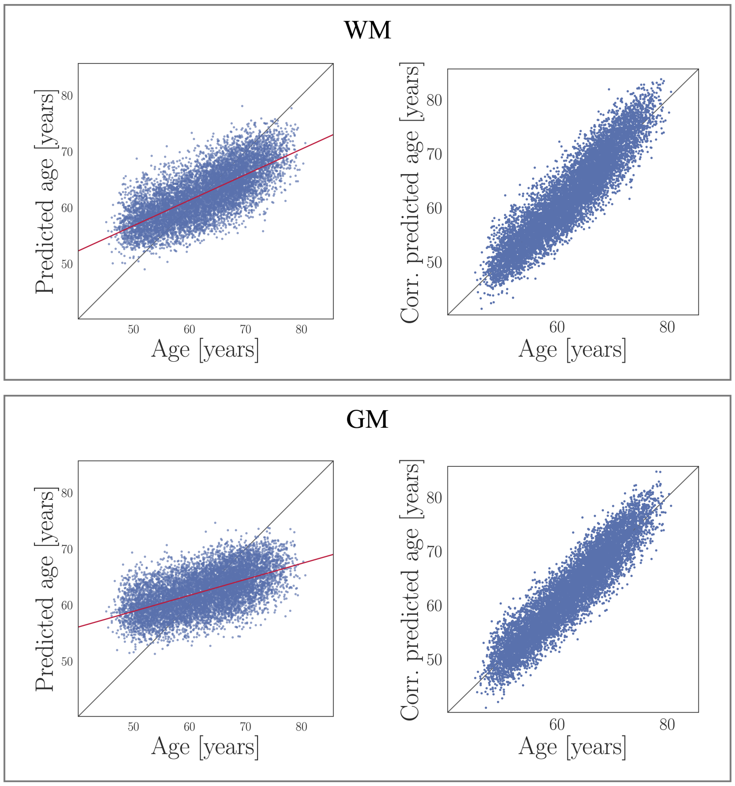


SI Figure 1: Age predictions for global white matter (WM) and grey matter (GM). The left plots show true age versus predicted age, with the red lines indicating the model fits ($r = 0.72[0.71, 0.73]$ for the WM model and $0.57[0.55, 0.58]$ for the GM model). The plots on the right show the predictions after correcting for age-bias [1, 2], which was done by first fitting $Y = \alpha \times \Omega + \beta$, where $Y$ is the modelled predicted age as a function of chronological age ($\Omega$), and $\alpha$ and $\beta$ represent the slope and intercept. The derived values of $\alpha$ and $\beta$ was then used to correct predicted age with $Corrected\ Predicted\ Age = Predicted\ Age + [\Omega - (\alpha \times \Omega + \beta)]$. This correction gives equivalent results to using age as a covariate in the linear regressions against number of previous childbirths [3].

#### 2. Correlations between brain age data, previous childbirths, demographics and covariates

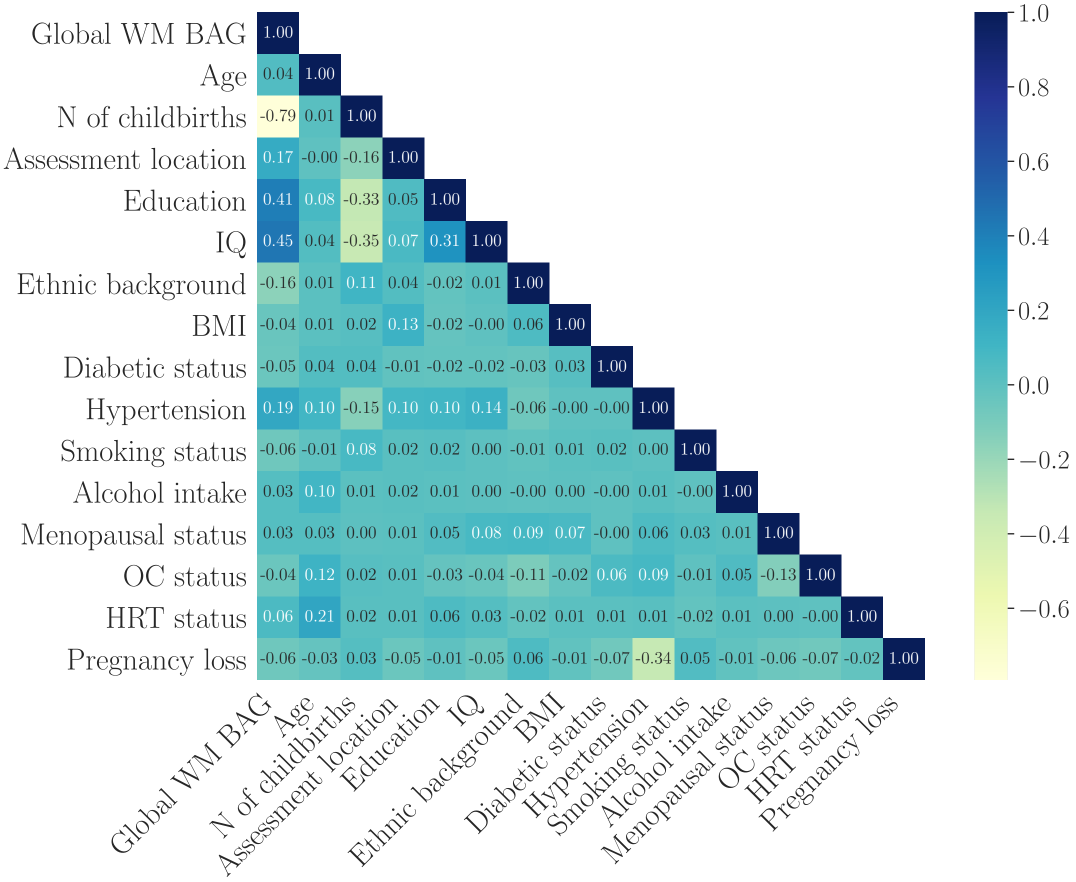


SI Figure 2: Correlations (Pearson's $r$) between global white matter (WM) brain-age gaps (BAG), number (N) of previous childbirths, demographics and covariates. IQ = intelligence quotient; BMI = body mass index; OC = oral contraceptive; HRT = hormone replacement therapy; pregnancy loss = experience with stillbirth, miscarriage or pregnancy termination.

#### 3. Associations with conventional MRI summary measures

SI Table 1: Associations between number of previous childbirths and the MRI summary measures total grey matter (GM) volume, mean fractional anosotropy (FA) and mean diffusivity (MD), as well as FA and MD in Corpus Callosum (CC) - the tract showing the strongest effects in the brain age analyses. Chronological age was included in the analyses for covariate purposes and $p$-values are reported before and after correction for multiple comparisons. All variables were standardised before entered into the analysis. SE = standard error.

| Measure | $\beta$ | SE | $t$ | $p$ | $p_{corr}$ |
|---|---|---|---|---|---|
| Total GM volume | 0.0204 | 0.010 | 2.009 | 0.045 | 0.064 |
| Mean FA | 0.0179 | 0.010 | 1.854 | 0.064 | 0.064 |
| Mean MD | −0.0262 | 0.010 | −2.675 | 0.007 | 0.022 |

#### 4. Feature importance ranking for brain age prediction in the corpus callosum

SI Table 2: Feature importance ranking for the Corpus callosum-based age prediction, with gain for each diffusion metric. The Gain indicates the relative contribution of the corresponding feature to the prediction model, calculated based on each feature's contribution for each tree in the model. A higher value implies that the feature was more important for generating the prediction. DTI = diffusion tensor imaging; DKI = diffusion kurtosis imaging; WMTI = white matter tract integrity; SMT = spherical mean technique; FA = fractional anisotropy; MD = mean diffusivity; AD = axial diffusivity; RD = radial diffusivity; MK = mean kurtosis, AK = axial kurtosis; RK = radial kurtosis; AWF = axonal water fraction; axEAD = extra-axonal axial diffusivity; radEAD = extra-axonal radial diffusivity; INVF = intra-neurite volume fraction; exMD = extra-neurite mean diffusivity; exRD = extra-neurite radial diffusivity.

| Feature | Gain |
|---|---|
| WMTI-radEAD | 2068.467539 |
| SMT-exMD | 1343.625053 |
| DTI-MD | 446.814543 |
| DTI-RD | 230.211839 |
| DTI-AD | 226.769257 |
| DTI-FA | 220.537127 |
| SMT-INVF | 213.289021 |
| DKI-AK | 210.312535 |
| SMT-exRD | 198.772824 |
| DKI-MK | 194.508129 |
| DKI-RK | 194.375870 |
| WMTI-AWF | 162.461406 |
| WMTI-axEAD | 157.201242 |
